# Simple Neurofeedback via Machine Learning: Challenges in real time multivariate assessment of meditation state

**DOI:** 10.1101/2022.09.27.509655

**Authors:** Sruthi Susan Kuriakose, Aishwarya Swamy, Rahul Venugopal, Arun Sasidharan

## Abstract

Attaining proficiency in meditation is difficult, especially without feedback since the mind may be easily distracted with thoughts and only long term efforts see any impact. Self-regulation would be much more effective if provided real time assessment and this can be achieved through EEG neurofeedback. Therefore, this work proposes a scheme for assessing meditation-like state in real time from short EEG segments, using low computational settings. Signal processing techniques are used to extract features from long term meditation practitioners’ multichannel EEG data. An autoencoder model is then trained on these features such that the model can be run in real time. Its reconstruction errors or its latent variables are used to provide non typical feedback parameters which are used to establish an objective measure of meditation ability. Our approach is optimised to have lightweight architectures handling small blocks of data and can be conveniently used on low density EEG acquisition systems as it requires only a few channels. However, our experimental results suggest that the meditation state has substantial overlap even in terms of multivariate EEG features and show prominent temporal dynamics, both of which are not captured using simple one class algorithms. Being an extremely flexible one-class model, we have described multiple improvements to the proposed autoencoder model to address the above issues and develop simple yet high precision neurofeedback protocols.

## I Introduction

The practice of meditation is often considered to help in attaining overall well-being. It is said to have therapeutic effects such as stress reduction, emotion regulation [1], as well as benefits such as increase in cognitive performance [2] and increased immune system activation [3]. It has also been shown to have positive impacts in people with anxiety, depression [4] and chronic pain [5]. Our goal is to assist individuals practice meditation since it appears to be the answer to a variety of medical problems associated with maladaptive mental health. Meditation takes prolonged focus, which is typically difficult for beginners compared to more seasoned practitioners. We address this challenge by proposing a real time neurofeedback framework designed to help novices in practicing meditation.

One way of quantifying the effects of meditation on the brain is using electroencephalograms (EEGs) which is a popular non-invasive technique used to record brain activity in clinical and research settings. At any given moment, ions flow in the neurons of the brain and groups of neurons produce synchronous activity, giving rise to voltage oscillations which can be measured using EEG devices [6]. EEG is characterised by its high temporal resolution, which allows for real time monitoring of signal changes. Since EEG signals are time series data, signal processing and machine learning techniques can be used to analyse and classify them in order to study distinct mental states. Real time approaches using EEG, also termed as Brain Computer Interfaces (BCIs) in the technology development domain and Neurofeedback in the clinical domain, have also been explored in various areas, particularly in emotion recognition [7] and epilepsy diagnosis [8]. These systems typically have 5 stages: (i) EEG signal acquisition stage, (ii) signal preprocessing stage (e.g., channel selection and band-pass filtering), (iii) feature representation learning stage, (iv) classifier training stage, and (v) feedback stage [9]. The feature extraction step often has a strong influence on the performance and sophistication of the classifier. It involves considerable domain expertise in order to analyse signals and optimise features [10], [11]. For classification, the features chosen for representation should be able to discriminate between the various brain states. Prior studies have proposed several methods to extract the various EEG features for different applications. For example, Chaudhary et al. [12] used spectral power features, statistical features as well as correlation features from the signal band power obtained from continuous Morlet wavelet transform in their work to detect primary colours from the features of raw EEG signals. Jana et al. [13] generated a spectrogram feature matrix from filtered EEG signals as an input to a one-dimensional convolutional neural network for seizure detection.

On the other hand, some researchers have chosen to entirely bypass the traditional signal processing techniques and use deep neural networks to perform the feature extraction from raw EEG data. They have taken the stance that advances in deep learning have reached the point where the model can automatically learn features that outperform human engineered features [14]. A two-dimensional supervised deep convolutional autoencoder has been able to detect epileptic seizures [11]. Similarly, another deep learning algorithm called Deep Motor features is found to be useful for a BCI with imagined motor tasks [15]. Li et al. [16] also used a variational autoencoder to determine the latent factors from multichannel EEG for the purpose of emotion recognition.

Although these techniques have good results, it would be a far simpler approach to devise the maximum possible feature set and then use an algorithm to learn the features which can best separate the classes. Too many features may decrease the efficiency of a real time system as well as accuracy of algorithms. We can then have a real time system which extracts only the necessary features and feeds it into a model to fit our goals. Our aim is to implement a computationally inexpensive algorithm in real time to assess how well an individual is meditating in real time and provide them with neurofeedback. The objective of this paper is not just to create a tool which can assess meditation ability, but also to create a framework involving machine learning methods which can be easily applied to any neurofeedback applications.

Currently, neurofeedback protocols which are popular in clinical applications focus on comparing the EEG features of patients with that collected from large number of normal individuals thereby creating a neurofeedback algorithm that rewards subjects (patients) whose EEG features approaching that of the general population [17]. Neurofeedback has been used in treatment of several mental disorders like ADHD, anxiety, depression, insomnia, epilepsy, stroke, addiction, autism, etc.,[18]. This study takes a similar approach wherein an algorithm is developed to help train users to make their EEG features similar to that of an expert practitioner in multiple meditation traditions. Users could perform a meditation practice and the neurofeedback device would non intrusively provide users with feedback on how well they are doing [19]. Currently, there are enterprises which use a similar setup of EEG devices and algorithms to train people to meditate via neurofeedback. However, the algorithms they use are proprietary, and it is unknown whether the data or features have sound scientific basis. As a result, we decided to develop an open source approach for evaluating meditation ability. The approaches mentioned by Brandmeyer and Delorme [19] in integrating neurofeedback and meditation were also instrumental in our work. Our objective is to build a model which can give us a real time feedback of our mental state, i.e., if it corresponds to meditative or non-meditative states. One major concern is that non-meditative states can be ambiguous and hard to define. While meditation may involve focusing attention, other activities such as being involved in a cognitively engaging task could also be considered as focusing attention. Hence, we may define non-meditative states as mind wandering during rest or even ruminating on negative thoughts. An ideal non-meditative state is hard to obtain, hence our proposed approach involves a one class algorithm, where non-meditating states can be determined by novelty detection.

Overall representation of this paper is presented as follows – Section-I contains an introduction with an overview of related works, Section-II describes the methods and experimental approaches which have been considered for this experiment, Section-III consists of the results and discussion on the methods, and Section-IV summarises the study and mentions future scope.

## II Materials and Methods

### A. Datasets and Preprocessing

In our study, we selected EEG data of meditators belonging to two different traditions - Brahma Kumaris Rajayoga meditators and Vipassana meditators (Buddhist mindfulness based practice). Both datasets were obtained from recordings done at the Centre for Consciousness Studies, Dept. of Neurophysiology, NIMHANS. All EEG data were acquired using a Geodesic EEG System 300 (Electrical Geodesics, Inc., USA) with NetAmps 300 amplifier and Net Station software version 4.5.7 and with 128 Channel HydroCel Sensornets. The signals were sampled at 1 KHz and channel locations were set as per the default channel file supplied by EGI.

#### 1) Rajayoga

The Rajayoga meditators had 48 participants belonging to a Long Term Practice group (LTP) who had a median of 14,240 h of meditation experience and a minimum of ten years of regular meditation practice [20]. The protocol design had two sets of rest and meditation states before and after a multi-level gamified cognitive task [21]. Each set contained rest eyes open (RO), rest eyes closed (RC), meditation eyes open (MO) and meditation eyes closed (MC) states. The MC state of the LTP group are chosen for the Meditating class. The Non-meditating class comprises both rest and cognitive task segments from both groups.

#### 2) Vipassana

The Vipassana meditators had two groups of practitioners - 22 Senior practitioners (those with over 7 years of daily practice along with atleast one long retreat) and 21 Teachers (instructors at meditation centres who have undergone several long retreats) [22]. Apart from the meditation EEG protocol, they were asked to do the same three-level sensory motor cognitive task mentioned above. For this study, we’ve selected 40 minutes of Vipassana data from their meditation EEG protocol for the Meditating class, as well as 4 minutes of eyes closed Rest condition and the cognitive task for the Non-meditating class.

#### 3) Preprocessing

In both the datasets, the signals were already preprocessed with band-pass filtering (between 1 and 40 Hz), artifact correction and removal (using the artifact subspace reconstruction (ASR) method) [35], bad channels interpolated (using spherical spline approach), down-sampling to 250 Hz, and re-referenced to M1 and M2 (mastoid electrodes). From each subject, four channels are chosen (F3, F4, P3, P4), as the bilateral frontal and parietal sites would represent a good part of the brain activity with the minimum electrodes and are present on most wearable EEG devices. These electrodes were chosen to reduce processing complexity and to have less setup time and maintain the subject’s convenience by having fewer electrodes. Moreover, the use of all channels may lead to an overfitting effect. The continuous component time series data was then epoched into 2 seconds with 0.5 second overlap for the Brahmakumaris dataset (since it was a smaller dataset) and 1 second overlap for the Vipassana dataset. A single EEG segment is represented as a matrix of shape (4 × L) where L is epoch length = sampling frequency (250) × segment duration (2). Equal number of segments were chosen from both the Brahmakumari and Vipassana groups. We made sure that the number of meditation segments (denoted by K) matched the number of non-meditating segments (500 segments each). Now, the EEG data for both groups is of dimensions (K × 4 × 500). Principal Component Analysis (PCA) is then applied on the data to reduce signal dimensionality and computation cost. PCA transforms the data into a new set of variables called principal components ranked according to variation explained in the original dataset. The first component with maximum variance is chosen as a channel. Any components with less than 50% variance are removed, as they may not be representative of the four-channel EEG segment.

### B. Feature Extraction

We are interested in finding the main signal processing features which could be used to distinguish between meditating and non-meditating EEG segments. Our processing pipeline is to first extract a number of features and then feed it into a machine learning model which learns the relative importance of features to classify the epochs. We could then extract the feature importance from the model by performing correlation analysis. Overall, we extract 60 features from the EEG data.

#### 1) Feature Set 1 - Spatial features (7)

When the PCA algorithm is applied for channel selection, we also get the relative weights of the channels for the principal component used. These weights represent spatial distribution or weights of the EEG component and are hence chosen as the main spatial features for extraction. The explained variance of that component is a measure of spatial coherence of the signal and is taken as another spatial feature. The right-left asymmetry in frontal and parietal weights and the average frontal-parietal asymmetry value are also calculated as additional spatial features.

#### 2) Feature Set 2 - Conventional spectral analysis features (13)

Spectral analysis is based on changes in specific rhythms and frequencies, which may be caused by physiological changes where neuronal populations synchronously oscillate at specific timescales [23]. We used Time-Frequency analysis to find the average power of each frequency band. Using Welch’s method, we compute a normalized value of total bandpower for all frequencies by scaling it down to an arbitrary upper limit of 100 microvolts, and relative bandpower for 7 bands - Delta (1-4 Hz), Theta (4-8 Hz), ThetaAlpha (6-10 Hz), Alpha (8-12 Hz), Beta1 (12-18 Hz), Beta2 (18-30 Hz), Gamma (30-40 Hz). We also evaluate ratios between different bands - alpha to theta, delta to theta, delta to alpha, alpha to beta1. We also compute an engagement metric according to the formula Engagement = Beta1 / (Alpha + Theta) [24].

#### 3) Feature Set 3 - Separating oscillatory and aperiodic features from spectral analysis data (31)

A drawback of spectral analysis is that it does not take into account the power law distribution; there is a 1/f background spectrum present under all conditions, even in minimally conscious states. Therefore, we want to analyse periodic features without bias from the aperiodic activity.

##### a) IRASA features (16)

The Irregular-Resampling Auto-Spectral Analysis (IRASA) method [25] analytically extracts the background spectrum (fractal/aperiodic component). In IRASA, the fractal component is identified by taking the median of geometric mean of power spectra of upsampled and downsampled signals. This is subtracted from the original power spectrum to get an estimate of the oscillatory power spectrum.

##### b) FOOOF features (15)

FOOOF (fitting oscillations & one over f) [26] is a tool similar to IRASA which tries to parameterize neural power spectra that were assumed to be the summation of 1/f (aperiodic component) and multiple narrowband oscillation humps (periodic component). The approach uses a precomputed power spectrum to iteratively apply Gaussian fits to all periodic components in order to obtain a model of the periodic part. This fit can then be removed from the spectrum to obtain an optimally pure aperiodic component.

Although both IRASA and FOOOF have the same goal of separating periodic and aperiodic data from the same spectrum, FOOOF removes the oscillatory component more parametrically than in IRASA, and hence the features captured have different values. Using both algorithms, we get the fit parameters (intercept, slope and R^2 of fitting an exponential function), and the oscillatory power spectral density (used to extract the same power spectrum features described above). IRASA also gives the standard deviation of the oscillatory component.

#### 4) Feature Set 4 - Non-linear features (8)

These are entropy and fractal dimension measures, which measure the irregularity and amount of chaos in a time series.

Entropy is often defined as the amount of uncertainty within a random variable. The system can be predicted with the greatest precision when the entropy is 0. [27]

##### a) Permutation entropy

it derives a probability distribution of the ordinal patterns by capturing the order relations between values in a time series [28].

##### b) Single Value Decomposition (SVD)

it measures the dimensionality of the data by giving a measure of eigenvectors required to adequately explain a data set.

##### c) Sample Entropy

it computes the conditional probability that two sequences similar for a set number of data points remain similar at the next point.

##### d) Lempel-Ziv complexity

It gives a measure of the number of different substrings encountered as the sequence is viewed from beginning to the end.

Fractal dimension (FD) of a signal is used to show its complexity and self similarity in the time domain [29]. Generally, higher self-similarity and complexity result in higher FD [30]. FD is expected to increase during calming meditation [31]. This can be interpreted as an increase in self-similarity of EEG signals during self-organization of hierarchical structure oscillators in the brain.

##### a) Detrended fluctuation analysis

it is used to find long-term statistical dependencies in time series. The data is split into multiple time windows, where each segment is detrended to avoid global and local trends.

##### b) Petrosian fractal dimension

it translates the series into a binary sequence by subtracting consecutive samples in the time series, and it computes a measure considering the number of sign changes in the binary sequence and the length of the sequence.

##### c) Katz fractal dimension

it is the ratio between the logarithms of the total length of the EEG time series and the normalized Euclidean distance between the first point to the point that provides the furthest distance.

##### d) Higuchi fractal dimension

it is based on a measure of length of the curve that represents the considered time series, while using a segment of k samples as a unit to measure the time series.

Entropy measures are highly affected by changing measurement scales, while fractal dimension measures are not. The fractal dimension thus captures a scaling symmetry independent of contraction or dilation, whereas entropy measures capture the spatio-temporal translational symmetry.

### C. Dataset Preparation

The above 60 features are extracted for each 2 second epoch from the meditation segments and the non-meditation segments. The EEG feature data for both groups is of dimensions (N x F) where N is no of segments and F is the number of features (F=60). Since the data included some ratios, where one of the values may be 0, the rows containing INF were dropped. Outlier data is then removed by using the Isolation Forest algorithm. The feature set is then scaled across each epoch to the [0,1] range using Min-Max normalization. This to ensure that the range of values in the original and the reconstructed segments are the same. The final number of segments after preprocessing is 20,230. We now take 80% of the meditating group for training, and 20% for testing. The testing dataset consists of an equal number of segments of meditating class and non-meditating class. The target column is now added for both groups, indicating 1 for meditation and 0 for non-meditation.

### D. Proposed System

The proposed system is a one class novelty detection algorithm which is trained on the extracted features from meditating EEG segments. We define a simple architecture involving an autoencoder model since an autoencoder model has the capability of learning the latent representation of meditative EEG features and can potentially detect when the subject is not meditating by measuring the anomalies in the feature data using the trained network. Autoencoders are a type of neural networks which can non-linearly compress the input into a lower-dimensional latent space and then reconstruct the output from this representation. As shown in Fig. 2., our model is a simple autoencoder where the encoder has 3 dense layers and the decoder also has 3 dense layers. It is trained with reconstruction loss set as mean average error and with Adam optimizer. The idea is to train the autoencoder with only the meditating class, and to capture the mean reconstruction error and set it as a threshold.

**Fig. 1.**
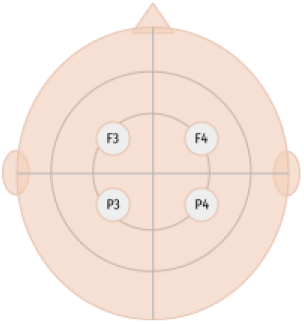
Topographical locations of sensors as per the International 10-20 System.

**Fig. 2.**
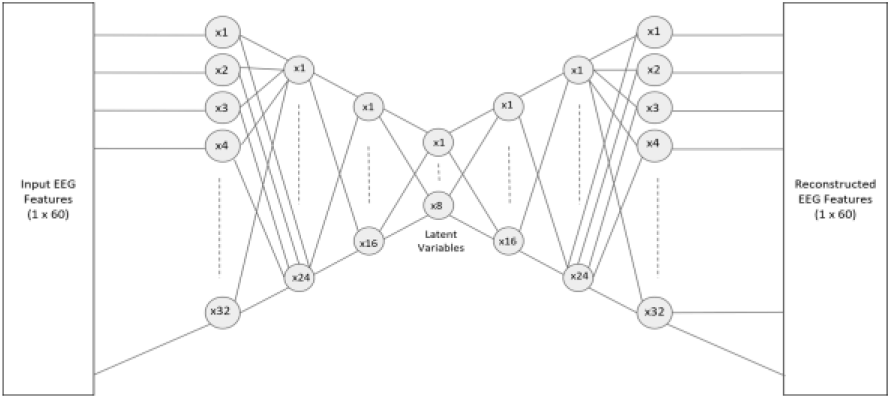
The architecture of the proposed autoencoder.

There are 3 approaches for classifying the meditation and non-meditation segments. One, the average reconstruction error of the training data is used as a threshold to obtain the classes. However, we also take into account that just taking the reconstruction error may seem too simplistic, hence we compute the mean reconstruction error for each of the 60 features, and set these as feature thresholds. We also calculate the feature count threshold as follows: Feature count threshold = mean number of features below the threshold + 1 standard deviation for the training data. The classification of the testing data is based on the number of features whose reconstruction error is below the threshold, and this is compared to the feature count threshold. Another method proposed for classifying the EEG features is to compare the latent representations. We calculate the latent variable thresholds as the range of the latent variable and follow the same approach above by calculating a threshold for the number of latent features whose values are in the meditation range.

#### 1) Comparison to other models

This model is compared to other one class algorithms such as One-Class Support Vector Machines (SVM) and Isolation Forest with default parameters. The one-class SVM [32] finds a hyper-plane in a multidimensional space that separates the given dataset from the origin, such that the hyperplane is as close to the data points as possible. A binary function is thus created that captures locations in the input space where the probability density of the data occurs. Isolation Forest [33] is a tree based algorithm based on isolated examples having a relatively short depth in the trees, while normal data is less isolated and has a greater depth in the trees.

#### 2) Model Implementation and Training

We implemented our models in Python programming language using many standard libraries, in particular, the Tensorflow machine learning library with its Keras deep learning API.

Both models are trained for 200 epochs (repetitions) on a batch size of 64, with an early stopping criterion of patience 5. The training data has a shape (no of samples * 1 classes, no of features) where no of samples = 16184.

### E. Evaluation metrics

The model is tested on two classes of meditating and non-meditating (rest and cognitive task) segments. The test data has a shape (no of samples x 2 classes, no of features) where no of samples = 4046. The evaluation metrics to assess classification efficiency of the models against the testing set include accuracy, precision, recall, and F1 score.

These evaluation metrics are defined as follows:

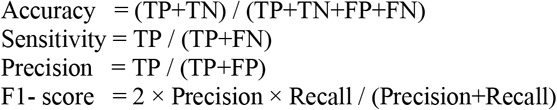

where P denotes the number of positive (meditating) EEG segments while N denotes the number of negative (rest or cognitive task) EEG segments. TP, TN, FP and FN are the numbers of true positives, true negatives, false positives and false negatives, respectively. Accuracy denotes the percentage of the correctly classified EEG features belonging to any state (meditating or non-meditating), sensitivity is the percentage of correctly classified meditating state features, while precision is the percentage of features classified as meditating state that are truly meditating state features. F1-score combines both precision and recall, such that the model with the highest score represents the model that is best able to classify observations into classes.

## III Experimental Results & Discussion

### A. Performance of the Model

From the results shown in Table I, it is apparent that none of the models seem to capture the differences between meditating and non-meditating EEG features well. The third autoencoder approach to create threshold ranges for latent variables gave very poor results as the range for the meditation latent features overlapped with that of the non-meditation latent features. This shows that one class algorithms cannot distinguish between these two states, i.e., that complex brain activity during meditation cannot be captured by using just the mainstream signal processing techniques. This approach needs further optimisations, which needs further understanding of the challenges.

**Table I:** Results of the 1-class machine learning models.

| Model | Accuracy | Precision | Recall / Sensitivity | F1-score |
| --- | --- | --- | --- | --- |
| Autoencoder (reconstruction error) | 0.54 | 0.52 | 0.90 | 0.66 |
| Autoencoder (feature count) | 0.49 | 0.50 | 0.82 | 0.62 |
| One class SVM | 0.48 | 0.49 | 0.81 | 0.61 |
| Isolation Forest | 0.53 | 0.52 | 0.87 | 0.65 |

### B. Feature overlap between states

Looking at Fig. 3, there is substantial overlap between the values of the 60 EEG features in meditating and non-meditating states. This could be an important reason for the failure of our proposed model as well as other one-class models to accurately classify these states.

**Fig. 3.**
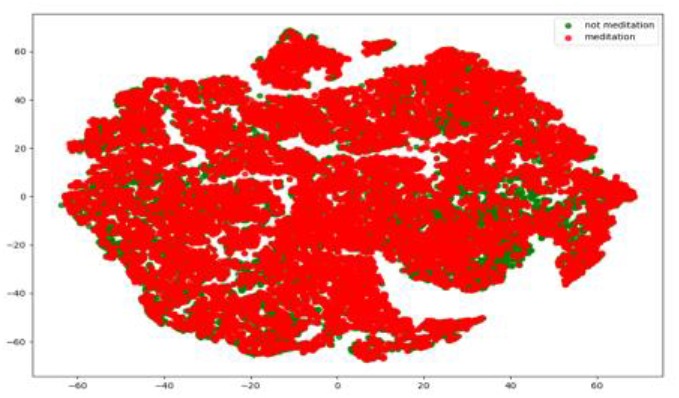
A high-dimensional visualisation showing the overlap between features for both classes captured by t-SNE.

Being a deep learning approach, the autoencoder models have the advantage over other single class models in that they can be optimised in numerous other ways. One such improvement we suggest is to create an ensemble of an autoencoder with a classifier such that they can be simultaneously trained [11]. This would ensure that the encoder will extract the latent information of the target class such that it also maximises its separation from other classes. However, the drawback of this model would be that besides having a large training dataset of our target condition (meditating state), we need similar amount of data for all possibilities of non-meditating states (which can include almost anything else including rumination on negative thoughts or daydreaming). We may also need to retrain the model each time we select a non-meditating state to avoid. Ideally, we would have our non-meditating state as a combination of the things we want to exclude. Another approach would be to create an ensemble of auto-encoders where each auto-encoder is associated with a unique class /state, making it more modular and not requiring retraining [34].

The main bottleneck in these models is identified to be the poor homogeneity among the different traditions of meditators. Although the primary focus of the paper had been to find common features within the state of meditation across traditions, it is evident that one class algorithms fail to capture such a signature. Hence, another approach is to follow the same pipeline taking only homogenous data. This approach was executed by training the autoencoder on the meditation features (46 features excluding the power ratios) of the Vipassana dataset and testing it on the rest and task features of the same. Here, we observe meditation F1-score of 0.83 and weighted F1-score of 0.8.

### C. Temporal dynamics of Features

None of the features consistently stay in a certain range of values in any of the states of rest, task or meditation. But, the dynamicity seems to show a trend for difference between, say Task and Meditation states. Hence, classifying every two seconds, without considering the temporal dynamics, as we have done in our approach, proves futile. A possible solution would be to use an algorithm that incorporates such temporal dynamics spanning across previous segments. This could be done by making an improvement to the autoencoder model, such as LSTM autoencoder. A convolutional autoencoder with features across time segments, may also be able to address the above issue.

From Fig. 4, we can see another important reason why the model does not show good evaluation metrics - brain states aren’t stable even during meditation. It could oscillate between the states, i.e., even during meditation, it may shift between features of meditation and rest. To extract the temporal dynamics of the data, 100 time domain features were added as downsampled time series data (to 25 Hz) giving a total of 110 features. However, this led to data scaling issues. Scaling the feature set within the sample decreased the performance of the model from a weighted F1-score of 0.5 to 0.39. In scaling across all the samples independently for each feature, the weighted F1-score is 0.44.

**Fig. 4.**
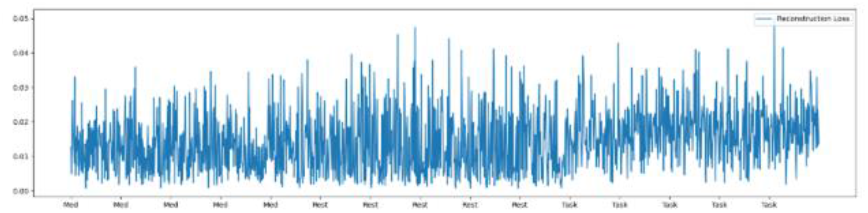
Reconstruction error of 500 consecutive epochs as a meditator transition across meditation, rest and task, using the proposed model.

The impact of scaling data on an autoencoder is further explored using a smaller dataset composed of 2 minutes of eyes open (EO) and eyes closed (EC) data which were epoched into 4 second segments from which several of the abovementioned features were extracted. The model trained on non-normalized data, data normalized within the sample, and data normalized across all the training samples gave a weighted F1-score of 0.47, 0.99 and 0.37 respectively. It was found that certain features which had its values from a higher range than the other features influenced the range of the normalized dataset in such a manner that the model was able to better differentiate between the two classes. However, when these features were removed from the dataset, the performance of the first two models decreased. So, scaling has several complex effects on the model performance and need to be evaluated well.

### D. Architecture of the Brain Computer Interface (BCI)

The autoencoder approach was taken, keeping in mind the need to provide a real time neurofeedback. Assuming the model is able to distinguish with some effectiveness between meditating and non-meditating data, the reconstruction error or the probability of the epoch as belonging to a meditation class can be given as neurofeedback in an auditory manner. This would be done with the help of subtle sound cues when their EEG features deviate from normative meditative features [19]. This real time approach creates a framework where subjective metadata can be fed as an input to help improve the model. Our end goal is to assist anyone practicing meditation in whether they’re moving in the right direction, hence we provide them with neurofeedback including metrics on how long they were able to meditate and with a meditation score. Our plan is to design a schema for a BCI system which can connect an EEG system and provide neurofeedback regarding meditation, as shown in Fig. 5. An EEG acquisition system has to be connected to Python, and sampling rate has to be set to 250 Hz. For example, the Neuroscan SynAmps RT EEG system can interface with MATLAB through its Curry software. The running MATLAB Session can be connected to Python. The Muse S can also be connected to Python through the BrainFlow python package. Once signals are obtained in real time, the EEG data of the four channels (F3, F4, P3, P4) is taken in 2 second windows and the 60 features are extracted from the signals. These features are given as an input to the model, and the model gives us a probability value on it being a meditation segment for neurofeedback. For each epoch, we provide an assessment of how well they’re meditating by adjusting the volume of an ambient noise when the probability value dips below the threshold to alert the user.

**Fig. 5.**
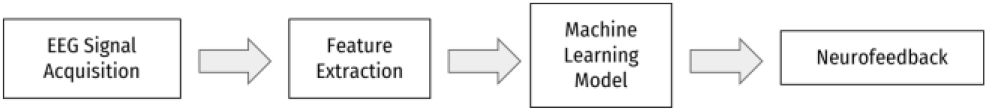
The Design Schema.

## IV Conclusions & Future Work

This paper explored EEG based outlier detection methods to produce a real-time multivariate assessment of meditation state. In this work, datasets from two different traditions of meditation are combined to extract multiple features which are then provided as an input to machine learning algorithms for the purpose of neurofeedback. This work attempts to bridge the gap between standard clinical neurofeedback protocols which are based on just 1 −2 features, and black box approaches used by companies that produce devices aimed at training people to have more focused and calmer minds. It is also an exploration of how autoencoders and other outlier detection models can be used for EEG data, as these kinds of subjective questions we seek to address such as how well someone is meditating tend to have unbalanced classes of just meditation data. Our approach is optimized to have lightweight architectures handling small blocks of data and can be conveniently used on low density EEG acquisition systems as it requires few channels. However, our experimental results suggest that the meditation state has substantial overlap even in terms of multivariate EEG features and show prominent temporal dynamics, both of which are not captured using simple one class algorithms. Being an extremely flexible one-class model, we have described multiple improvements to the proposed autoencoder model to address the above issues. Besides, we plan to examine if use of a convolutional autoencoder to directly extract features from raw data, can address the issues. However, these models require larger amounts of data and they tend to be less interpretable as the models are often very complex and sophisticated. We are also interested in checking if the model can perform better on shorter or longer EEG segments. Considering the variability in brain activity features within meditation sessions, we believe that real-time ratings from subjects during a neurofeedback session using our pre-trained models, could help further optimise the precision of the algorithm, at each individuallevel.

## Acknowledgment

We acknowledge Dr Ajay Kumar Nair and Dr Ratna Jyothi, whose PhD dataset at the Center were used for this study. We thank the other investigators of the project (Dr Bindu M Kutty, Dr Ravindra PN, Dr Geetha Desai and Dr Ramajayam G), CCS and Axxonet team for their support.

